# Transcriptomic comparison of *Drosophila* snRNP biogenesis mutants: implications for Spinal Muscular Atrophy

**DOI:** 10.1101/044693

**Authors:** Eric L. Garcia, Ying Wen, Kavita Praveen, A. Gregory Matera

**Affiliations:** Integrative Program for Biological and Genome Sciences; Lineberger Comprehensive Cancer Center; Departments of Biology and Genetics, University of North Carolina at Chapel Hill, Chapel Hill, North Carolina, 27599, USA

**Author notes:** Current address: Regeneron Genetics Center, Tarrytown, NY 10591, USA.

**Keywords:** alternative splicing, alternative polyadenylation, alphaCOP, Abelson interacting protein, dUTPase, GARS, Survival motor neuron, SMN, Spinal muscular atrophy, SMA, Phosphorylated adaptor for RNA export, PHAX, Ars2, Prp6, Prp8, RNA-sequencing, snRNA, snRNP biogenesis

## Abstract

Spinal Muscular Atrophy (SMA) is caused by deletion or mutation of the *Survival Motor Neuron 1* gene (*SMN1*), but the mechanism whereby reduced levels of SMN protein lead to disease is unknown. SMN functions in the assembly of spliceosomal small nuclear ribonucleoproteins (snRNPs) and potential splicing defects have been uncovered in various animal models of SMA. We used disruptions in *Smn* and two additional snRNP biogenesis genes, *Phax* and *Ars2*, to classify RNA processing differences as snRNP-dependent or *Smn* gene specific in *Drosophila*. Although more numerous, the processing changes in *Ars2* mutants were generally distinct from those identified in *Phax* and *Smn* animals. *Phax* and *Smn* null mutants exhibited comparable reductions in steady-state snRNA levels, and direct comparison of their transcriptomes uncovered a shared set of alternative splicing changes. Transgenic expression of *Phax* and *Smn* in the respective mutant backgrounds significantly rescued both snRNA levels as well as alternative splicing. When compared to the *Smn* wild-type rescue line, three additional disease models (bearing SMA-causing point mutations in *Smn*) displayed only small-to-indistinguishable differences in snRNA levels and the identified splicing disruptions. Comparison of these intermediate SMA models revealed fewer than 10% shared splicing differences. Instead, the three *Smn* point mutants displayed common increases in stress responsive transcripts that correlated with phenotypic severity. These findings uncouple organismal viability defects from the general housekeeping function of SMN and suggest that SMN-specific changes in gene expression may be important for understanding how loss of SMN ultimately causes disease.

## Introduction

Spinal Muscular Atrophy (SMA) results from genetic loss of the human *Survival Motor Neuron* 1 (*SMN1*) gene [1], but uncovering how SMN protein depletion leads to the characteristic neuromuscular pathology of SMA has proven difficult. SMN functions in the assembly of Sm-class small nuclear ribonucleoproteins (snRNPs), core components of the spliceosome [2,3]. Because of this well-established function, prior studies investigating SMA etiology have sought to connect SMN depletion to alternative splicing changes and downstream SMA symptoms [4-6]. However, these studies typically do not account for indirect effects, like developmental delays, that might be responsible for the observed differences [7,8]. Thus, the evidence linking a reduction in snRNP levels to spliceosomal dysfunction and specific splicing changes is confounded by false positives. Hence, whether or not SMA phenotypes are caused by disruptions to snRNP levels and pre-mRNA splicing remains unsettled.

*Drosophila* models of SMA exhibit numerous small changes in splicing patterns that are difficult to separate from biological noise and normal developmental fluctuations [7]. In the fruit fly, steady-state levels of small nuclear RNAs (snRNAs) appear to be uncoupled from SMA-like phenotypes [9]. Studies in SMA mouse models also failed to uncover clear connections between SMN depletion, snRNP-levels, and pre-mRNA splicing. Although late-stage symptomatic mice show widespread splicing changes [4], these differences are not apparent prior to the onset of symptoms [10]. In these studies, global alternative splicing patterns may have masked important cell-type specific differences. Recently, tissue-specific disruptions to the splicing patterns of genes in motor neurons and glial cells were found in pre-symptomatic SMA model mice [6]. However, it is not clear that decreased snRNP levels caused the observed changes to the splicing pattern. Thus, disruptions to snRNPs and splicing in motor neurons and glial cells of the central nervous system (CNS) may not be the sole drivers of SMA pathology.

Evidence from fruit fly and mouse SMA models suggests that high SMN levels are required in cells and tissues outside of the CNS. A combination of pan-neuronal and mesodermal specific SMN expression is required to rescue lethality in *Drosophila* SMN mutants, as expression in either cell-type alone was insufficient for rescue [11]. Similarly, restoring full SMN expression solely within motor neurons of SMA mice does not rescue viability [12,13]. Furthermore, SMA-like phenotypes in severe mouse models that utilize a human *SMN2* transgene can be rescued by peripheral administration of splice-correcting antisense oligonucleotides (ASOs) [14]. Intriguingly, restoration of SMN expression in the CNS was not essential for phenotypic rescue, as CNS-specific treatment with a blocking oligonucleotide complementary to the splice-correcting ASO did not hamper the rescue [15]. Additional work in fly SMA models has shown that the entire neuronal circuit, including proprioceptive interneurons, is important for motor function, indicating that SMN plays a non cell-autonomous role in the maintenance of neuromuscular connections [16]. Thus, SMA is not a cell-autonomous defect of motor neurons in mice and flies, and studies that focus solely on RNA changes in motor neurons would thereby neglect important changes in other cell types.

Although the evidence connecting SMN loss (and rescue) to particular splicing changes is unclear, observations in diverse eukaryotes suggest that perturbations of the core splicing machinery can affect specific alternative-splicing events. Indeed, mutations in yeast genes encoding core spliceosomal factors disrupted the splicing of only a subset of pre-mRNAs [17,18]. Inhibiting highly expressed ribosomal protein coding genes in yeast also perturbed numerous alternative splicing patterns, suggesting that spliceosomal accessibility controls pre-mRNA splicing patterns [19]. In *Drosophila*, core splicing factors were uncovered by an RNAi screen for regulators of a set of alternative splicing reporters [20]. In cultured human cells, knockdown of Sm proteins led to numerous changes in alternative splicing [21]. Thus, evidence from multiple systems indicates that modulating snRNP levels can alter splice site choice, however, a clear connection between snRNP levels and human disease has yet to be made.

In this study, we carried out transcriptome profling of *Drosophila Smn* mutants, along with two additional snRNP biogenesis mutants in order to identify snRNP-dependent vs. SMN-specific changes in gene expression, premature cleavage and polyadenylation, and alternative splicing. Specifically, mutations in genes encoding the snRNA-specific nuclear export adaptor Phax (<u>Ph</u>osphorylated <u>a</u>daptor for RNA e<u>x</u>port) and the Cap Binding Complex (CBC) associated protein, Ars2 (<u>Ars</u>enite resistance <u>2</u>), were used for comparison [22-28]. The *Phax* mutant was particularly useful in identifying an overlapping set of snRNP-dependent RNA changes in our SMA model flies. We found that alternative-splicing differences shared between the *Phax* and *Smn* mutants can be rescued at both the RNA and protein levels by expression of *Phax* or *Smn* transgenes, respectively. In combination with a modest restoration of steady-state snRNA levels, observations from these rescue lines provide a connection between snRNP levels and specific pre-mRNA splicing events. However, the link between these changes and SMA phenotypes remains unclear.

## Results

### *Phax* and *Smn* mutants display similar steady-state snRNA decreases

RNA-sequencing (RNA-seq) of poly(A)-selected RNA was used to identify mRNA changes in snRNP biogenesis mutants, and northern analysis coupled with RNA-seq of total ribosomal RNA-subtracted (rRNA-) RNA was used to quantify steady-state snRNA levels. Browser shots of *Phax, Smn*, and *Ars2* gene loci confirmed the gene disruptions in the respective mutant lines (Fig. 1A; [7]. In our previous RNA-seq analysis of *Smn* null mutants, publicly available modENCODE data were used to determine the developmental stage at which the mutants arrested [7]. By the same method, the *Phax* and *Ars2* mutants exhibited a similar timing of developmental delay as did the *Smn* animals (Fig. S1A&B).

**Figure 1.**
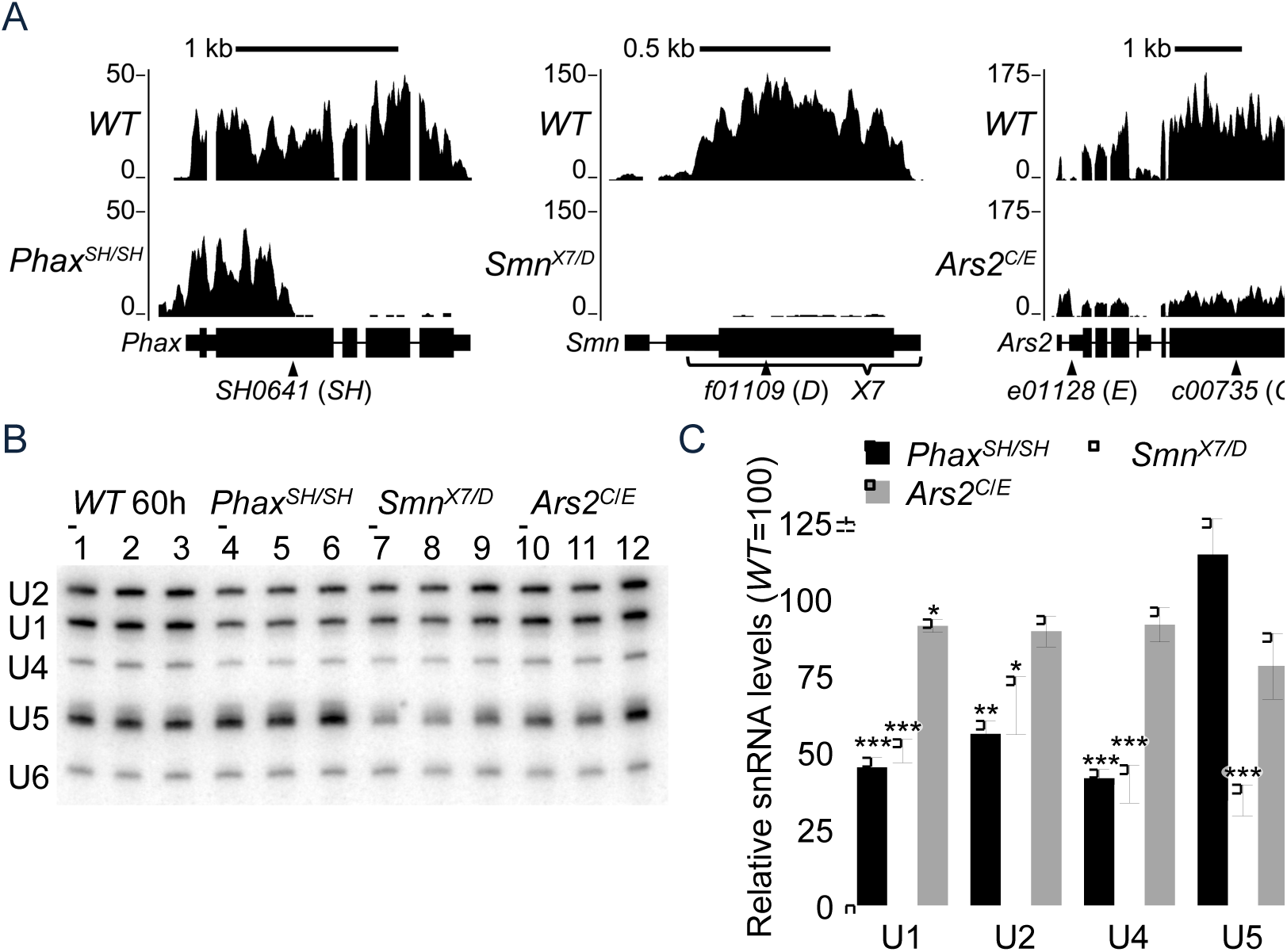
Analysis of three different snRNP biogenesis mutants. (A) Browser shots of larval RNA-seq reads at the three disrupted gene loci. Reads from *Oregon-R* (*WT*) versus *Phax*^*SH/SH*^*, Smn*^*X7/D*^, or *Ars2*^*c/E*^ snRNP biogenesis mutants are shown. Mapped read tracks were normalized to the median of the middle two quartiles of the mapped *WT* sequence read counts. (B) Northern blot of RNA from snRNP biogenesis mutants and *WT* larvae. RNA was extracted from *WT* larvae at 60 ± 2 hours (60h) post egg-laying, and mutant RNA was extracted at 74 ± 2 hours to account for their delayed development (see text). (C) Quantification of northern blot in (B). Sm-class snRNAs were normalized to the Lsm-class U6 snRNA, and U6-normalized *WT* snRNA levels were set at 100. Asterisks are *p*-values from a Student’s t-test: * *p*-value < 0.05, ** *p*-value < 0.01, and *** *p*-value < 0.001.

Using developmentally appropriate wild-type controls (Fig. S2A), northern blots demonstrated comparable, near two-fold decreases in steady-state U1, U2, and U4 snRNAs in the *Phax* and *Smn* mutants (Fig. 1B&C). When assayed by northern blotting, U5 snRNA levels were essentially unaffected in the *Phax* mutants (Fig 1B&C), as were levels of nearly all of the snRNAs in the *Ars2* mutants (Fig. 1B&C). When RNA-seq data were used to quantify snRNA levels, each of the three mutants appeared to exhibit snRNA deficits, but the extent varied by the method of normalization (Figure S2B&C). Only small (<15%) changes were observed in specific snRNA paralogs relative to the total (Fig. S3), quantified by using reads covering isoform-specific nucleotides [29]. Overall, the *Phax* and *Smn* mutants displayed the most consistent decreases in steady-state snRNA levels.

### snRNP biogenesis mutants share a small number of gene expression changes

To identify shared and distinct changes in gene expression between snRNP biogenesis mutants, we used the TopHat/Cufflinks pipeline to quantify mRNA differences from our poly(A)-selected RNA-seq data [30]. Raw reads from age-matched wild-type (*WT*) controls and modENCODE second (L2) and early third instar (L3 12h) developmental controls were integrated with sequencing data from the mutants into a single analytical pipeline. For comparison, we focused on genes whose expression changed uniformly and significantly (false discovery rate-adjusted *p*-value < 0.05) relative to each of the controls (L2, L3 12h, and our *WT*). A relatively small number of genes and undifferentiated loci with multiple genes (186 out of ~15,000) met these stringent criteria for differential expression (i.e. 186 loci were significantly different from all of the controls; total number from Table S1), and most of these differences were mutant specific (Fig. 2A&C and Tables S1-S4). A number of stress-responsive transcripts were among those that were specifically elevated in *Smn* mutants, though some of these were also up slightly in the *Ars2* mutants (Fig. 2B). *Phax* mutants shared only twenty-eight gene expression changes with *Smn* mutants (Fig. 2C and Table S2). This small list of genes was enriched for genes involved in oxidation-reduction, as measured by DAVID gene ontology term analysis (*p*-value = 0.003) [31-33]. A notable gene expression change shared by the *Phax* and *Smn* mutants was a 5-fold increase in the *Activity-regulated cytoskeleton associated protein 1* (*Arc1*) transcript (Table S2). Arc1 functions in a well conserved hyperlocomotion response to starvation, and hence elevated levels in *Phax* and *Smn* mutants might be an indirect consequence of a disruption in nutrient acquisition in these animals [34]. In summary, gene expression analysis uncovered a small number of overlapping changes that exceeded normal developmental fluctuations. The relationship of these changes to SMA etiology and/or pathology remains to be determined.

**Figure 2.**
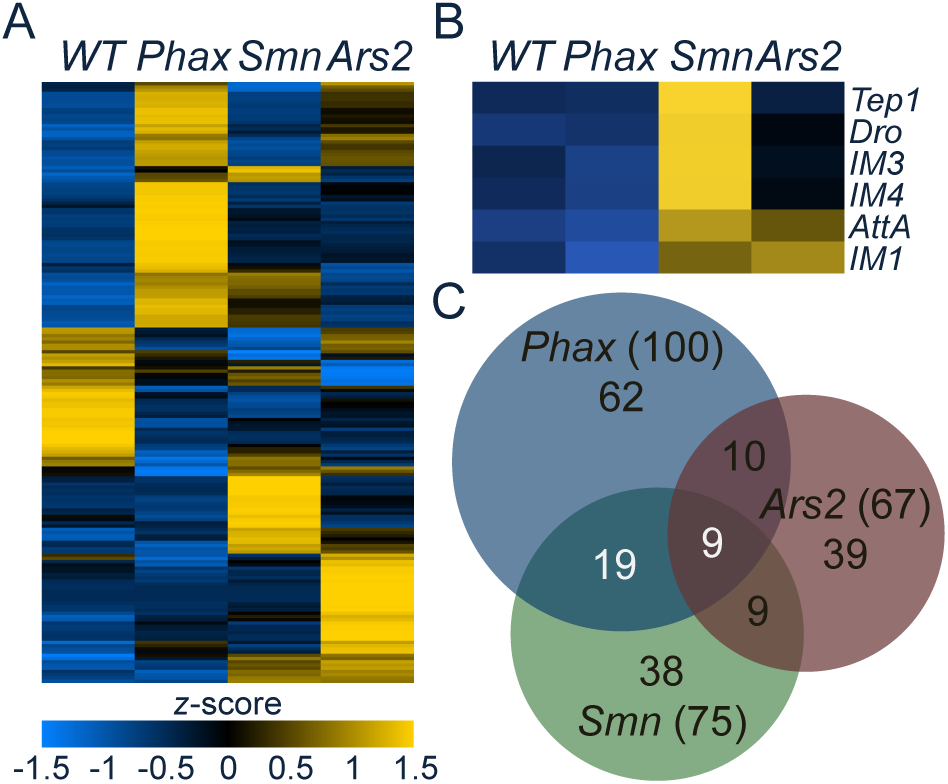
Gene expression differences in snRNP biogenesis mutants. (A) Heatmap comparison of Cuffdiff FPKM levels of differentially expressed transcripts. Heatmap colors were rescaled for each row and the rows were clustered based on pattern of gene expression between *WT* and mutants. (B) A heatmap of FPKMs from a set of stress-responsive transcripts in (A). (C) Venn diagram of the overlap in gene expression differences in the snRNP biogenesis mutants. *Phax* = *Phax*^*SH/SH*^, *Smn* = *Smn*^*X7/D*^, and *Ars2* = *Ars2*^*C/E*^. Numbers in parentheses are totals.

### snRNP biogenesis mutants exhibit a trend toward shorter mRNAs

In addition to its role in splicing, U1 snRNP has been reported to control mRNA length by preventing premature cleavage and polyadenylation [35,36]. To determine whether reduced U1 snRNA levels in our snRNP biogenesis mutants correlated with expression of shorter mRNAs, mapped RNA-seq reads were analyzed using the DaPars linear regression algorithm [37,38]. DaPars output is displayed as a Percentage of Distal PolyA site Usage Index (PDUI) for each transcript, or as the difference in the PDUI value between wild-type and mutant (ΔPDUI). Pairwise comparisons of PDUI values for wild-type and mutants can be used to visualize changes in RNA length. Whereas the *Phax* and *Smn* mutants expressed transcripts that were both longer and shorter than their wild-type counterparts, there were greater numbers of shortened mRNAs (Fig. 3A-C and Tables S5-S7). Focusing on differences that are greater than two-fold, *Phax* and *Smn* mutants had a relatively small number of mRNA length changes in common with each other (34). However, the changes were largely distinct from those observed in *Ars2* animals, who had more than twice as many mRNA length changes as did the *Phax* and *Smn* mutants combined (Fig. 3D and Table S8). The individual direction of length change, either longer or shorter, was also factored into the consideration of overlapping differences in Fig. 3D. The overall trend of shorter transcripts in the mutants does not appear to be a consequence of developmental delay, as parallel DaPars analyses of modENCODE datasets of second to third instar larvae showed a trend toward shorter transcripts in the more mature animals (Fig. S4). Among the few overlapping mRNA length changes between *Phax* and *Smn*, one of the largest and most apparent changes was a shortened mRNA for *CG9662*, a putative oligosaccharyl transferase complex subunit (ΔPDUI = -0.96, Table S5). These results are generally consistent with the notion that reductions in U1 snRNP levels predispose mRNAs to premature cleavage and polyadenyation. Because the *Ars2* mutants exhibited only a small decrease in steady state U1 levels (<10%; Fig. 1C), the mRNA length changes in these animals are more likely due to disruptions of the CBC, which are known to affect a subset of pre-mRNA termination events [26,39,40].

**Figure 3.**
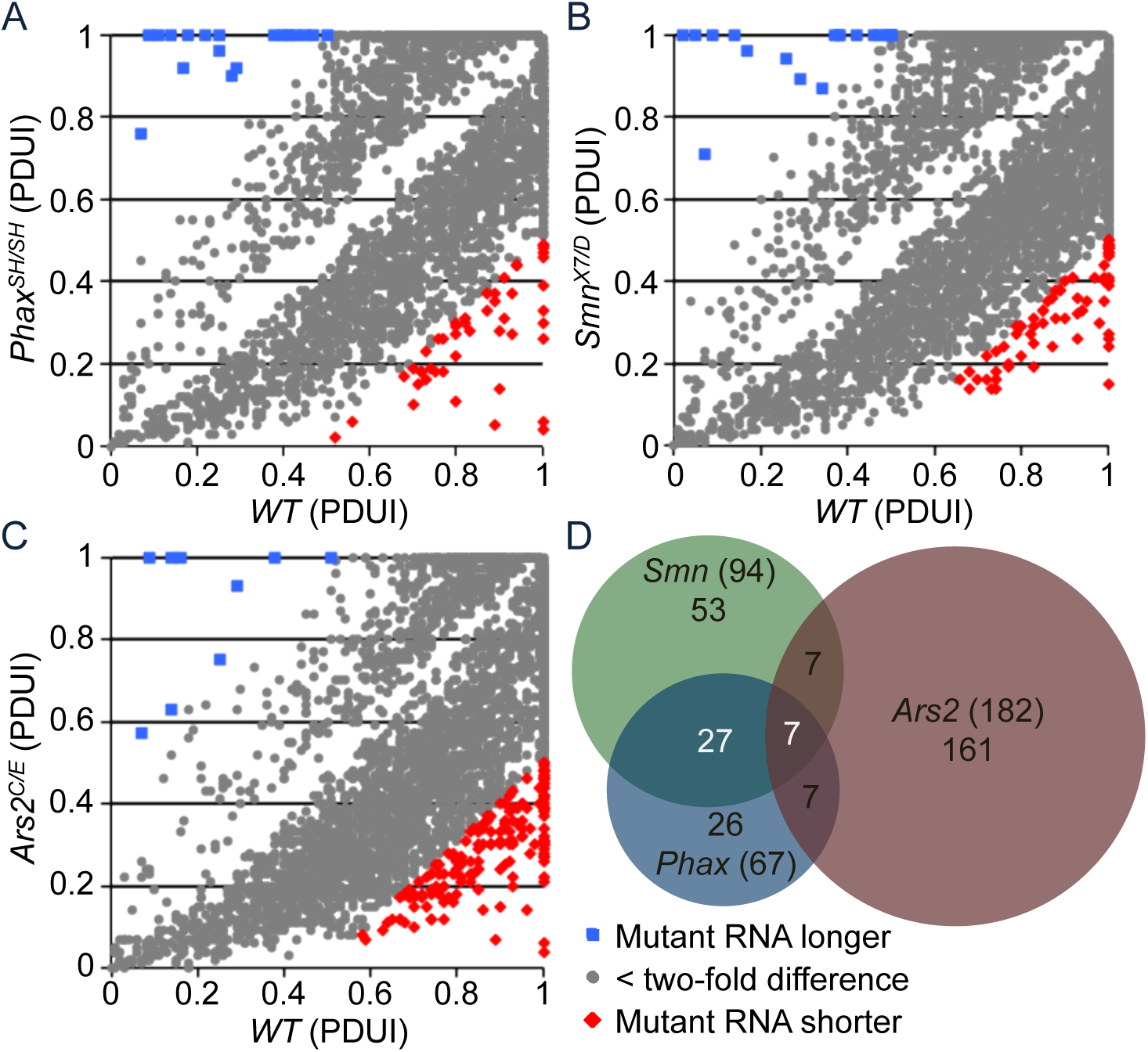
mRNA length changes in snRNP biogenesis mutants. Pairwise comparison of significant (False discovery rate adjusted *p*-value < 0.05) differences in mRNA length between: (A) *WT* and *Phax*^*SH/SH*^; (B) *WT* and *Smn*^*X7/D*^; and (C) *WT* and *Ars2*^*C/E*^. RNA length differences were extracted from mapped RNA-seq read files by the DaPars linear regression algorithm (see text). Percentage of Distal PolyA site Usage Index = PDUI. (D) Venn diagram of RNA length differences in snRNP biogenesis mutants labeled as in Fig. 2.

### *Phax* and *Smn* mutants share a set of alternative splicing pattern changes

Our previous RNA-seq analysis of *Smn* null animals revealed many small-amplitude alternative splicing differences (compared to wild-type controls), but it was unclear whether any of these changes might be due to decreases in snRNP levels [7]. To ascertain whether any of these splicing changes are snRNP-dependent, we used the Mixture of Isoforms (MISO) probabilistic framework to compare splicing differences in *Smn* mutants with those identified in *Phax* and *Ars2* mutants [41]. Simply put, the MISO algorithm estimates the expression of alternatively spliced isoforms in RNA-seq data, and reports it as a fraction of alternatively spliced events relative to the total [41]. MISO denotes this fraction as the ‘percent spliced in’ or ‘Psi’ [41]. Differences are reported here as an absolute change relative to the wild-type control, |deltaPsi|. Similar to our previous results with the *Smn* null animals, we found mostly small changes (less than two-fold, |deltaPsi| < 0.5) in alternative-splicing events in the *Phax* and *Ars2* (Fig. 4A-C and Tables S9-S11). More than half of the splicing changes identified in the *Smn* null animals overlapped with changes in the *Phax* mutants, but neither the *Phax* nor *Smn* mutants had considerable overlap with those observed in the *Ars2* mutants (Fig. 4D and Table S11). Of the top 150 differences in the *Ars2* background, close to ninety percent of these changes were located in the 5'-proximal intron (Table S11). These changes likely reflect the association of Ars2 with the CBC and the role of this complex in promoter proximal splicing events [24] rather than Ars2-related changes in snRNPs. In contrast, the changes that were common to *Phax* and *Smn* mutants are more likely to be due to decreases in snRNP levels.

**Figure 4.**
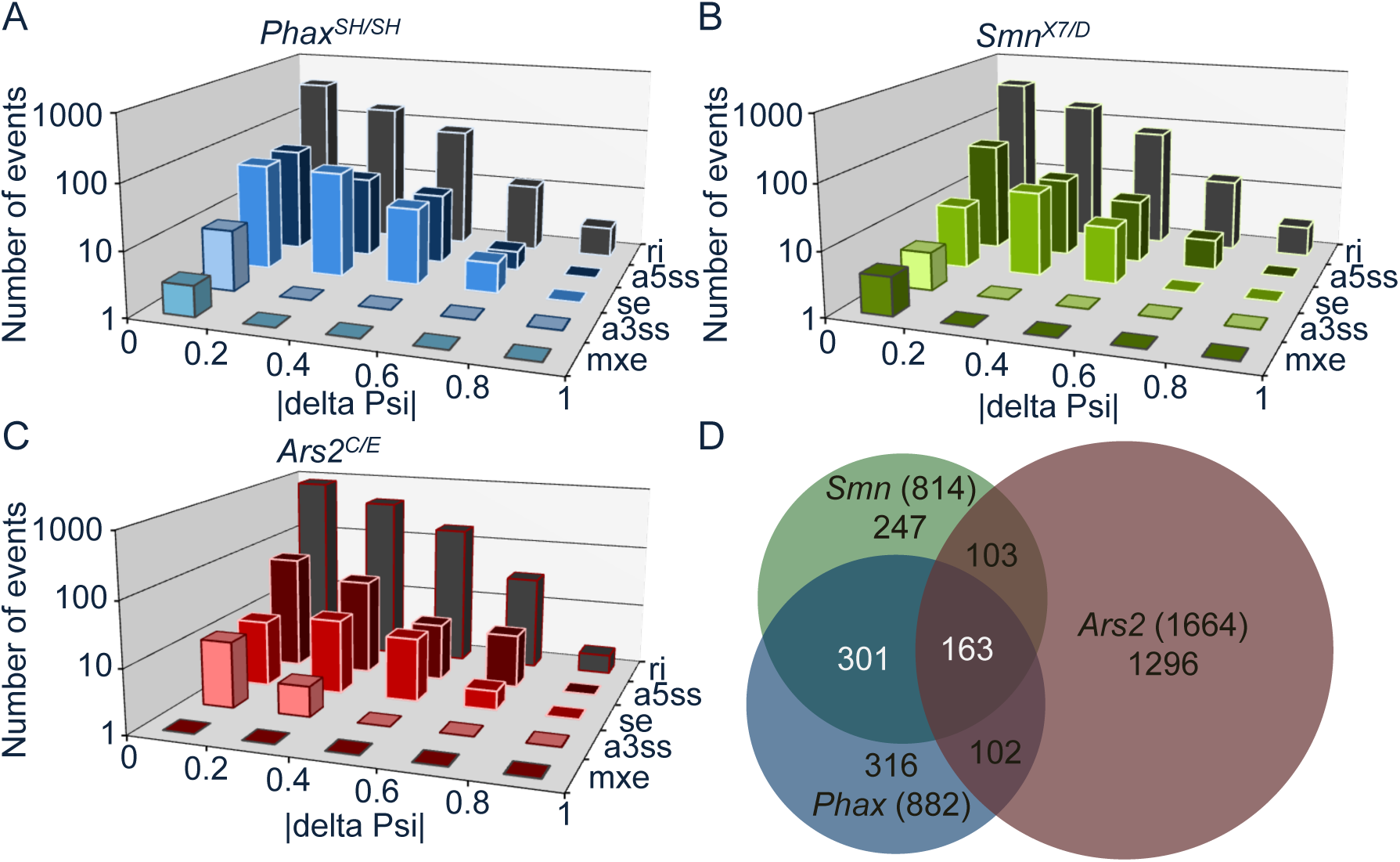
Alternative-splicing differences among snRNP biogenesis mutants. Distribution of significant alternative-splicing changes from small |delta Psi| near 0 to large |delta Psi| near 1: (A) *Phax*^*SH/SH*^ mutants relative to WT; (B) *Smn*^*X7/D*^ mutants relative to WT; and (C) *Ars2*^*C/E*^ mutants relative to *WT*. (D) Venn diagram of overlapping alternative-splicing changes in snRNP biogenesis mutants, labeled as in Fig. 2.

### Transgenic rescue of alternative splicing changes induced by loss of Phax or Smn

As described previously, *Smn* is essential for viability in the fly [11], and transgenic expression of Flag- or GFP-tagged SMN protein in an otherwise null mutant background fully rescues the larval lethality and other SMA-like phenotypes [42,43]. Here, we found that *Phax* is also an essential gene, and that ectopic expression of *Phax-GFP* using the GAL4-UAS system rescued the lethality of *Phax*^*SH/SH*^ mutants (Table S12). We next assayed (and confirmed by semi-quantitative RT-PCR) a number of the most apparent alternative splicing changes identified by RNA-seq (Fig. S5). We then tested the ability of the *Phax* and *Smn* transgenes to rescue four of these splicing events by quantitative qRT-PCR (Fig. 5B&C). Ubiquitous expression of *Phax* using an *Armadillo-GAL4* driver significantly rescued all of the splicing changes we tested (Fig. 5B). Transgenic expression of *Smn*, driven by its native promoter, also rescued these alternative-splicing events in *Smn* null animals (Fig. 5C). Together, the results confirm that the observed changes we identified by RNA-seq are indeed caused by loss of *Phax* or *Smn* expression.

**Figure 5.**
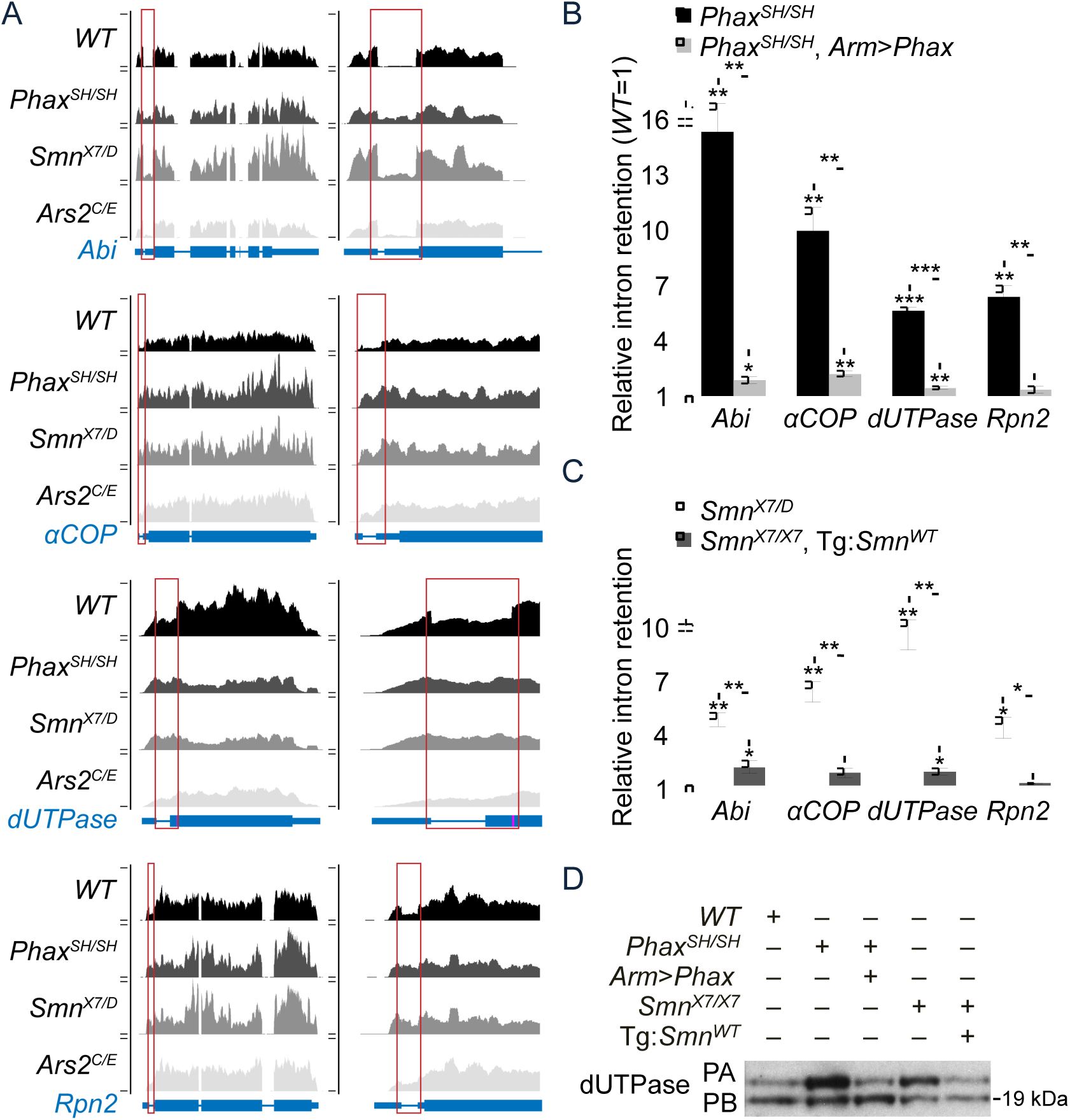
Transgenic rescue of alternative-splicing changes in *Phax* and *Smn* mutants. (A) UCSC browser shots of mapped read tracks for four of the top fifteen identified alternative splicing changes. (B) qRT-PCR analysis of intron retention in *Phax*^*SH/SH*^ mutants versus *Phax*^*SH/SH*^ mutants expressing a *UAS-Phax* transgene from the armadillo promoter driven GAL4 (*Arm>Phax)*. (C) qRT-PCR of intron retention in *Smn*^*X7/D*^ mutants versus *Smn*^*X7/X7*^ mutants expressing a wild-type *Smn* transgene from its native promoter. For (B) and (C), levels of *WT* intron retention were set at 1. (D) Western blot for dUTPase. Alternate protein isoforms reflect alternative-splicing change in *dUTPase* mRNA. Wild-type *Smn* transgene in *Smn*^*X7/X7*^ background = Tg:*Smn*^*WT*^. Long dUTPase protein isoform = PA and short = PB.

One of the splicing changes we detected is expected to produce an alternative (larger) dUTPase protein isoform. Therefore, we assayed the impact of this alternative splicing change by western blotting, using an antibody that targets both dUTPase isoforms. As shown in Fig. 5D, the expected increase in the ratio of the long isoform relative to the short one was observed in the mutants, and this change was largely rescued in both transgenic lines. Thus, the transcriptomic changes we identified in SMA model animals can (and do) translate into detectable differences at the protein level.

An increase in steady-state snRNA levels correlates with rescue of alternative splicing If the alternative splicing changes, observed in both *Phax* and *Smn* mutants, are truly caused by the decreased availability of snRNPs, then rescue of steady-state snRNA levels should mirror the rescue of these alternative splicing changes in the *Phax* and *Smn* transgenic lines. Northern blotting was used to determine whether the expression of *Phax* and *Smn* transgenes increased steady-state snRNA levels in the mutants (Fig. S6A&B). Expression of the *Phax* transgene by the *Armadillo* driver significantly rescued U1 and U4 snRNA levels in the *Phax*^*SH/SH*^ mutant background (Fig S6A&B). All of the snRNAs tested here were significantly rescued by expression the wild-type *Smn* transgene in the *Smn* null background (Fig. S6C&D). For the wild-type *Smn* rescue line, these observations differ from our previously published results [9], wherein we detected a modest increase in snRNA levels in the rescue line. This difference likely reflects minor improvements to our northern blotting protocol, specifically: using less total RNA per gel lane and more labeled probe in the hybridization reaction to ensure probe excess. The increases in steady-state snRNA levels that we observed in the *Phax* and *Smn* rescue lines correlated well with the phenotypic rescue at both the organismal (viability) and molecular (alternative splicing) levels.

### Splicing changes shared between *Phax* and *Smn* mutants are also caused by knockdown of splicing factors Prp6 and Prp8

Although the correlation of snRNA levels with particular splicing events provides evidence that these events are acutely sensitive to disruption of the core spliceosomal machinery, it does not rule out other possibilities. For example, the covariance could be due to a direct interaction of splicing factors with Phax and SMN, or a result of indirect factors, such as altered developmental progression. To test whether the observed splicing changes are indeed linked to a reduction in snRNPs and a corresponding disruption in spliceosome function, we depleted the tri-snRNP and U5-associated proteins Prp6 and Prp8 using RNA interference. As shown in Fig. S7, SMN depletion caused a slight (<two-fold) perturbation in the splicing events tested, but the Prp6 and Prp8 knockdowns disrupted the splicing of these transcripts to a much greater extent (Fig. S7A-C). Knockdown of Prp8 disrupted the splicing of the events identified above (*Abi*, *αCOP*, *dUTPase*, and *Rpn2* intron 1 (int. 1)), but it did not significantly disrupt the splicing of a downstream intron in *Rpn2*, which serves here as a control for broader spliceosomal dysfunction. The *Prp6* and *Prp8* knockdowns also altered dUTPase protein expression patterns to a greater extent than did knockdown of *Smn* (Fig. S7B). Steady-state snRNA levels were not significantly affected by any of these knockdowns, with the possible exception of U5 snRNA, which was slightly reduced (~40%) in the Prp8 depletion (Fig. S7D&E). The lack of snRNA decreases in the SMN depleted samples is consistent with the well-known perdurance of snRNPs in cultured cells, and it likely mirrors the observed low-amplitude disruptions to splicing. Prp6 and Prp8 are core spliceosomal proteins, and their depletion is expected to severely compromise the spliceosome [44,45]. Thus, the splicing changes seen in these knockdowns are perhaps not unexpected, but they nevertheless support the conclusion that the splicing changes identified in the *Phax* and *Smn* null mutants are snRNP dependent.

### Hypomorphic *Smn* missense mutants rescue snRNP levels and do not display defects in alternative splicing

As mentioned, the snRNP biogenesis mutants described above are severe loss-of-function alleles. SMA is, however, a hypomorphic condition, as complete loss of SMN activity is lethal in all organisms tested. To better model the disease in the fly, we previously engineered an allelic series of transgenic animals that express *Smn* missense mutations known to cause SMA in humans [9,46]. In addition to the wild-type rescue transgene (*Smn*^*WT*^), we analyzed three hypomorphic lines that display intermediate (SMA type II) phenotypes: *Smn*^*T205|*^*, Smn*^*Y107C*^, and *Smn*^*V72G*^ [46]. These lines cover a range of phenotypic outcomes: *Smn*^*T2051*^ is semiviable (~30% eclose as adults), *Smn*^*Y107C*^ is pharate-lethal (very few animals eclose), and *Smn*^*V72G*^ mutants all die as pupae [46]. Importantly, in this system, the wild-type and mutant transgenes are integrated at the identical chromosomal locus and are expressed from the native promoter in an otherwise *Smn* null background [9,46]. Using qRT-PCR, we analyzed the alternative splicing patterns of the four transcripts described above. As shown in Fig. 6A, none of the *Smn* missense lines exhibited a greater than two-fold change (in either direction) relative to *Smn*^*WT*^. Consistent with this finding, the missense mutants also displayed little to no difference in steady state snRNA levels, compared to the wild-type rescue line (Figs. 6B&C). To ensure that differences in developmental timing did not confound the analysis, animals were harvested after they began to display the wandering behavior that is characteristic of the late third instar stage of *Drosophila* development. We also measured snRNA levels of animals at slightly earlier and later developmental time points, and we found little difference between the mutant and the control lines (Fig. S8). Steady-state snRNA differences in earlier animals may reflect subtle differences in developmental progression from wild-type controls, but these decreased levels were largely absent from the later pupal stage, which only exhibited significantly lower U4 levels in the single *Smn*^*Y107C*^ line.

**Figure 6.**
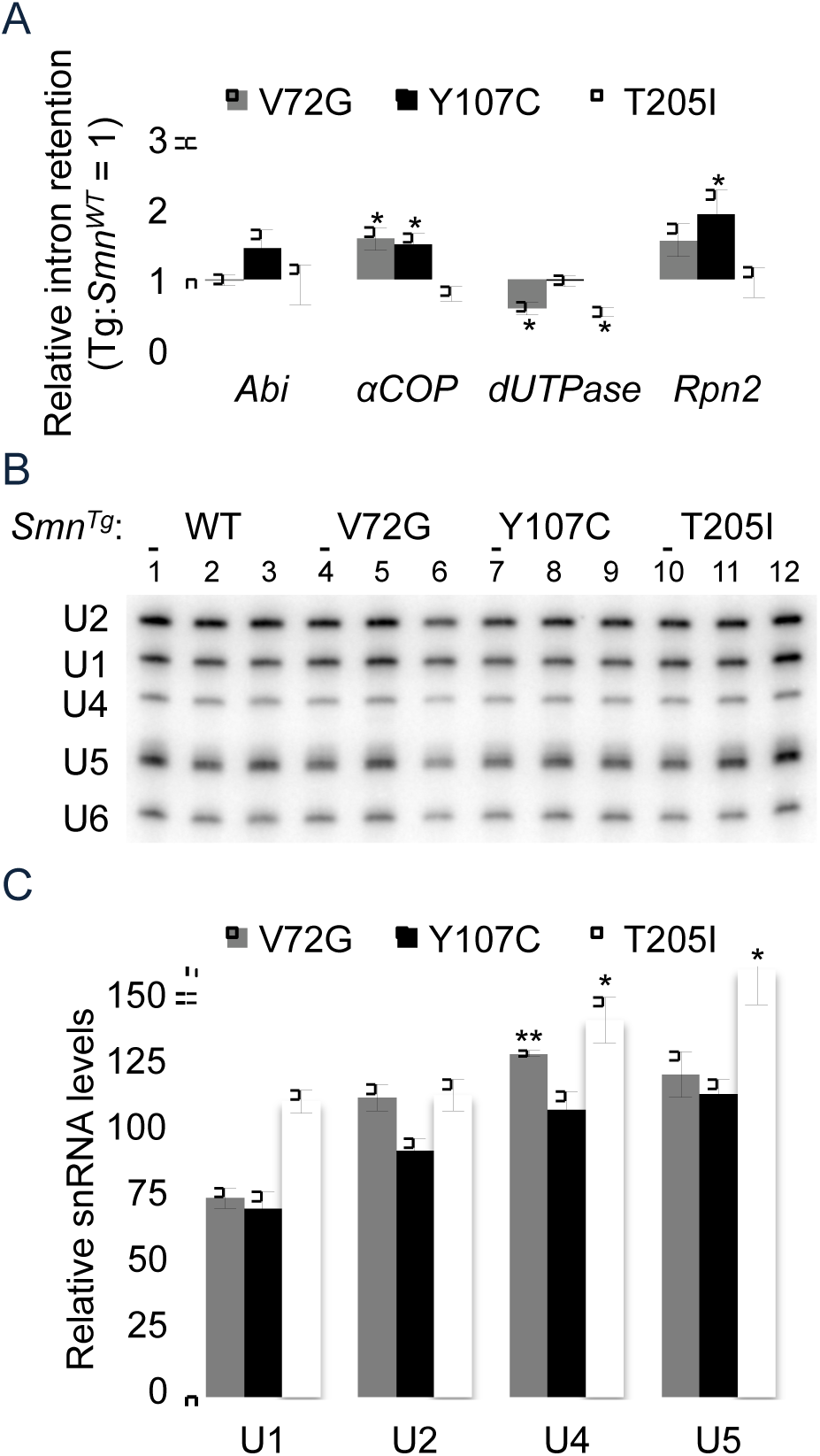
Analysis of steady-state snRNA levels and alternative splicing of target genes in *SMN* missense mutants (A) qRT-PCR analysis of intron retention of flies expressing SMA patient-derived missense mutations. Transgenic animals of the following generalized genotype were used: *Smn*^*X7/X7*^*,Flag-Smn*^*Tg/-*^, where Tg represents a WT, V72G, Y107C, or T205I transgene. Intron retention in the *Smn*^*WT*^ transgenic rescue line was set at 1. (B) Northern blots of snRNA levels in the missense mutant liness. (C) Quantification of snRNA levels from panel B. RNA levels in the *Smn*^*WT*^ (WT) transgenic line were set at 100.

### *Smn* missense mutants display limited overlap in splicing pattern changes

The large-amplitude changes identified in the severe loss-of-function mutants were not evident in the *Smn* hypomorphs, however these lines could exhibit other splicing pattern or gene expression changes that might account for differences in their phenotypic outcomes. To identify RNA changes in the hypomorphic lines, we sequenced total (rRNA-subtracted) RNA from whole animals isolated ~12hr after puparium formation, a developmental stage that *Smn* null animals never reach. The hypomorphs displayed numerous changes in their splicing patterns relative to the wild-type rescue line (1662 combined; Tables S13-16). However, the missense mutants displayed few large amplitude (greater than two-fold) changes (107/1662), and hardly any of these events were shared between them (42/1662; Fig. 7A and Table S13). Of the forty-two overlapping splicing changes, only six were previously identified as being snRNP-dependent (four) or *Smn* gene-specific (two) by comparison to data from the severe *Phax* and *Smn* loss-of-function lines (Fig. 7B). Browser shots of representative examples are shown in Fig 7C. We used qRT-PCR to validate one of the two *Smn* gene-specific changes. The *scarface* transcript (Fig. S9 and Table S13) exhibited an increase in intron retention between the third and fourth exons in the *Smn*^*V72G*^ transgenic line (deltaPsi = -0.61 and a >100-fold change by qRT-PCR). In the fly, Scarface functions as negative regulator of JNK (c-Jun N-terminal kinase) stress signaling [47]. As measured by RNA-seq, the *Smn*^*Y107C*^ (deltaPsi = -0.23) and *Smn*^*T205|*^ (deltaPsi = -.07) mutants exhibited smaller changes; these events were below the limit of detection by qRT-PCR (Fig. S9A and TableS13). Consistent with the notion that this is not a snRNP-dependent change, an increase in *scarface* intron retention was not observed following knockdown of Prp8 (Fig. S9B). Thus, RNA-seq profiling of hypomorphic, SMA-causing missense mutations is in agreement with the qRT-PCR data described above, and together these findings demonstrate that the splicing changes identified in the *Smn* null animals are not conserved among the intermediate SMA models.

**Figure 7.**
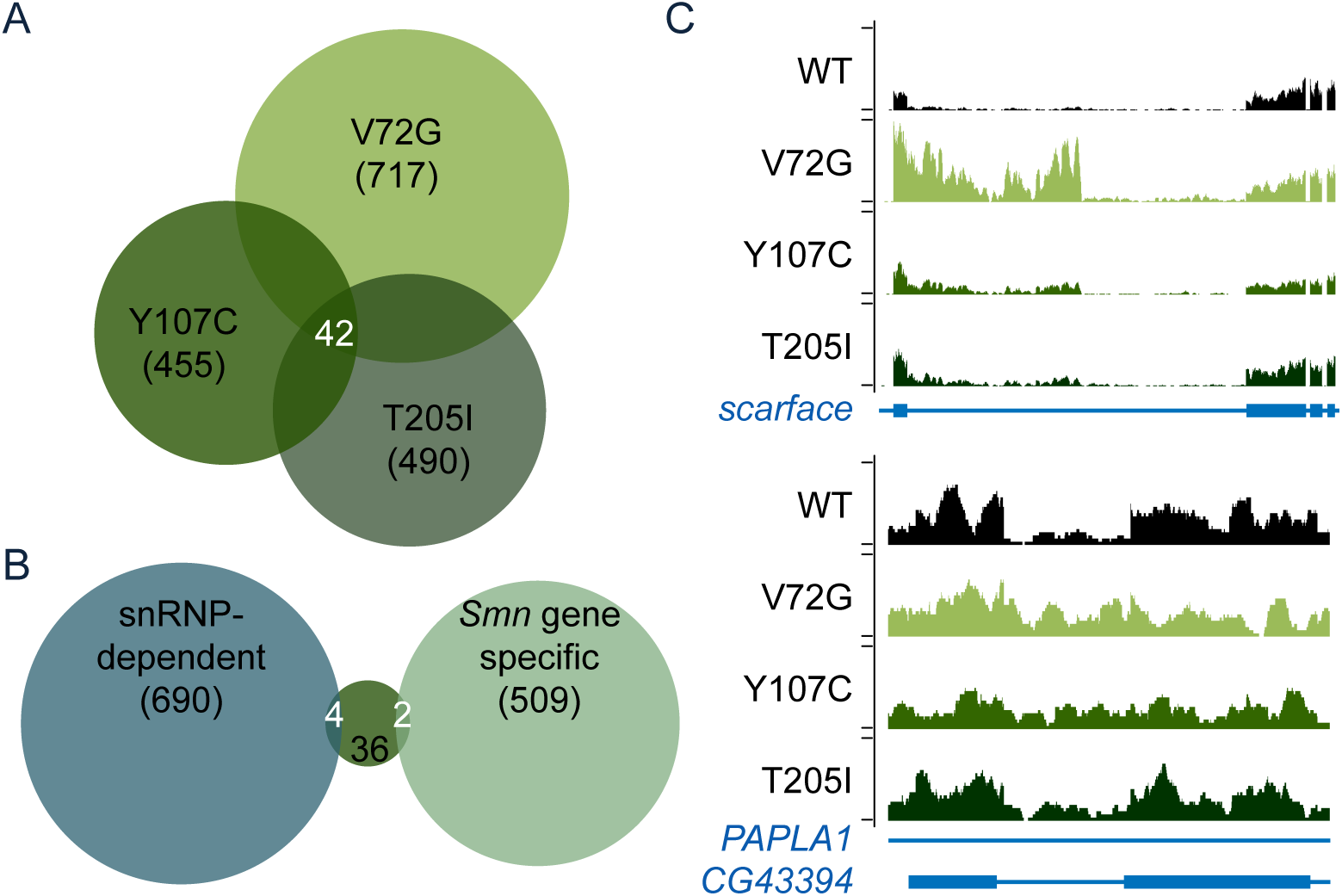
Alternative splicing changes in Smn missense mutants. (A) Venn diagram of overlapping alternative-splicing changes in *Smn* missense mutants. (B) Venn diagram of the overlap in shared splicing differences in the *Smn* missense mutants with either snRNP-dependent changes or *Smn* gene-specific changes categorized by comparison of the *Phax* and *Smn* total rRNA(-) RNA-seq. (C) Browser shot examples of an *Smn* gene-specific change in alternative-splicing in *scarface* (top) or a snRNP-dependent change in *CG43394* within a *PAPLA1* intron on the opposite strand (bottom). Genotypes are labeled by *Smn*^*Tg*^, which was expressed in the *Smn*^*X7*^ null background.

### Differential expression analysis reveals an activation of stress signaling in *Smn* missense mutant animals

In addition to changes in splicing patterns, our genome-wide analysis of RNA from the severe loss-of-function mutants revealed potential snRNP-dependent and *Smn* gene-specific changes in steadystate mRNA levels. Differential expression analysis of the *Smn* missense mutant lines revealed numerous changes in mRNA abundance in the most severe line, *Smn*^*V72G*^ (599), but far fewer in the other less severe lines, *Smn*^*Y107C*^ (88) and *Smn*^*T2051*^ (59) (Fig. 8A&B and Table S17-S21). Despite the low number of changes in the least severe *Smn*^*T2051*^ line, more than half of these overlapped with the other hypomorphic mutants (Fig. 8B). Surprisingly, many of the stress-responsive transcripts identified in the severe *Smn* null mutants were also among the mRNAs that were upregulated in all three *Smn* transgenic lines, relative to the wild-type rescue line (Fig. 8C and Table S18). As corresponding increases in mRNA levels were not found in the *Phax* mutants (Fig 2B), these events were classified as *Smn* gene-specific. Thus, unlike the snRNP-dependent splicing changes (which are not conserved in the *Smn* hypomorphs), the intermediate models of SMA exhibited clear RNA signatures of stress that correlate with disease severity (Fig. 8C) and are independent of decreases in snRNP supply (Fig. 2B).

**Figure 8.**
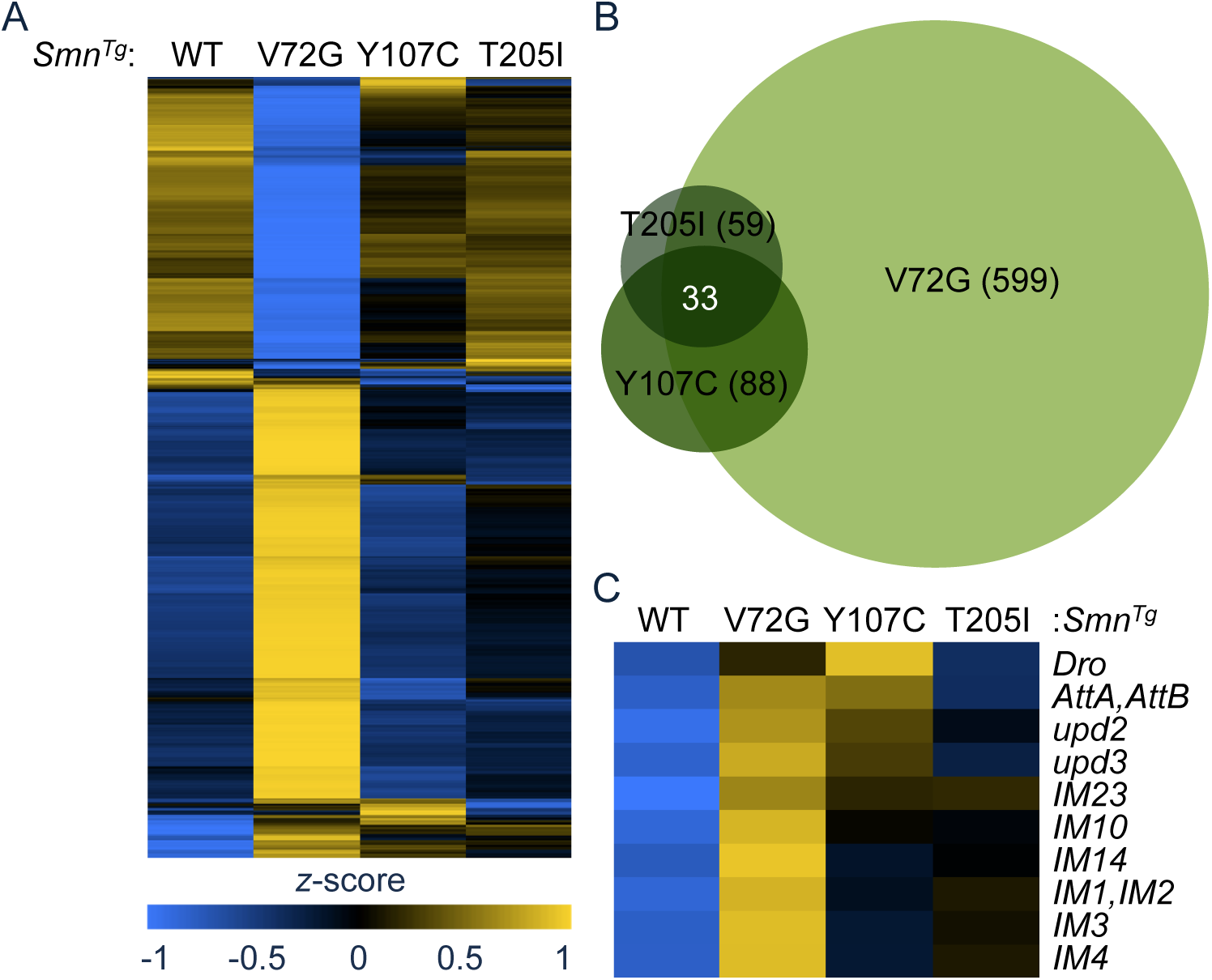
Differential gene expression in *Smn* hypomorphic animals. (A) Heatmap comparison of Cuffdiff FPKM levels of differentially expressed transcripts. Heatmap colors were rescaled for each row and the rows were clustered based on pattern of gene expression between *Smn*^*WT*^ rescue line and the *Smn* missense mutants. (B) Venn diagram of overlapping differences in mRNA levels. (C) A heatmap of FPKMs from a set of stress-responsive transcripts. Genotypes are labeled by *Smn*^*Tg*^, which was expressed in the *Smn*^*X7*^ null background.

## Discussion

Connecting SMA pathology to SMN’s function in snRNP assembly and downstream expression of specific mRNA isoforms remains a necessary benchmark for defining how splicing changes might contribute to disease etiology. Therefore, a first step toward uncovering the molecular mechanisms that elicit SMA outcomes would be to separate snRNP-dependent phenotypes from those that are SMN specific. Transcriptomic profiling of *Smn* and *Phax* mutants enabled us to identify a robust set of snRNP-dependent alternative splicing changes in SMA model flies. We show that reductions in steady-state snRNA levels correlate with the appearance of these alternative-splicing events, and that rescue of snRNA levels in transgenic animals correlates with the restoration of normal splicing. Furthermore, depletion of core splicing factors (Prp8 or Prp6) essentially bypasses the snRNP depletion step, causing these same alternative-splicing events, and providing a more direct link to spliceosomal dysfunction. Together, these observations support a close connection between snRNP levels and proper gene expression *in vivo*. Whether or not disruptions to this ubiquitously required process contribute to SMA phenotypes is less clear.

Ars2, 5' exons, and U1 telescripting Changes in splicing and mRNA length in *Phax* and *Smn* animals were largely distinct from the more abundant changes in the *Ars2* mutants (Figs. 3D and 4D). Ars2 is a component of the 5' Cap Binding Complex (CBC), and this association may underlie the transcriptomic changes that we identified in this mutant [48,49]. The *Ars2* splicing changes were mostly in 5'-proximal introns, and this finding is consistent with CBC mutants in yeast, *Xenopus*, and plants [50-56]. The association of Ars2 with the CBC likely also underlies the numerous mRNA shortening events we observed. These findings are consistent with previous studies in yeast and human cells that suggest the CBC protects a subset of weak poly(A)-signals from premature cleavage and termination [26,39,40]. Importantly, both types of Ars2 dependent mRNA processing events were largely distinct from the snRNP-dependent changes observed in the *Phax* and *Smn* mutants.

The distinct mRNA processing changes in *Ars2* mutants may also reflect species-specific differences in the role of the CBC and Ars2. Unlike *ARS2* knockdown experiments in mammalian cells [26,57], we did not observe defects in snRNA or histone mRNA processing. This finding is consistent with the observation that Ars2 protein is dispensable for proper histone mRNA maturation in *Drosophila* cells [58], and it suggests that aspects of Ars2 function are not conserved between vertebrates and invertebrates.

Our identification of a large number of mRNA shortening events that are shared between *Phax* and *Smn* mutants supports a recently discovered role for the U1 snRNP in the protection of premRNAs from premature cleavage and polyadenylation (PCPA). PCPA was previously observed by subtractive hybridization PCR, coupled with high throughput sequencing of RNA from cells treated with antisense oligomers targeting the 5'-end of U1 snRNA [35,36]. This protective role for U1 snRNPs was termed ‘telescripting’ [35,36] and this study is the first to validate this process outside of cell culture models. Although we observed an overall trend toward shorter transcripts in our datasets (Fig. 3), we found only a few large-magnitude (greater than two-fold) changes (Table S2). This is perhaps unsurprising, given that treatment of cultured cells with relatively high concentrations of anti-U1 oligomers provides an acute and nearly complete inhibition of U1 snRNA function, whereas the mutations in *Phax* and *Smn* do not perturb U1 to the same extent in the mutant animals.

RNA signatures of disease Alternative splicing depends on the regulated accessibility of pre-mRNAs to the spliceosome. Studies in a variety of eukaryotes have shown that changing this accessibility, either directly or indirectly, alters the splicing patterns of specific pre-mRNAs [17-21]. The extent to which snRNP levels contribute to normal splicing regulation and/or disease pathology is still being worked out, however, changes in the pattern of alternative-splicing have been reported in various SMA model systems [4-6,10]. The splicing changes reported in these SMA models are typically assumed to stem directly from loss of SMN and snRNPs, but such changes are often difficult to distinguish from broad, indirect effects of disease pathology or developmental delay [7,10]. By identifying overlapping splicing changes in *Phax* and *Smn* mutants, we are now able to distinguish between snRNP-dependent changes and gene-specific effects.

Many of the alternative splicing changes shared by *Phax* and *Smn* occur in genes of known importance to neuromuscular biology. For example, the mammalian ortholog of the alphaCOP protein has a demonstrated interaction with SMN, and it co-localizes with SMN in neuronal projections [59,60]. This interaction was recently shown to be required for neurite outgrowth in mammalian cell and zebrafish models of SMA [61]. Orthologs of the Abi protein appear to have important roles in neurogenesis via the regulation of actin dynamics [62-64]. Dominant mutations in GARS cause the distal neuropathy Charcot-Marie-Tooth disease type 2D, and recent observations suggest that GARS may colocalize with SMN in human cells [65-67]. Mouse models of SMA exhibit disruptions to ubiquitin homeostasis, and evidence of SMN interactions with proteasomal components, including Psmd1, the mouse ortholog of *Drosophila* Rpn2 [68,69].

Despite our finding of splicing disruptions in genes of known importance to SMA and neuromuscular biology in severe loss-of-function mutants, these splicing changes were absent from less severe models of SMA. Splicing patterns are not conserved from flies to humans, and we would not expect to find similar splicing disruptions in mammalian models of SMA. Finding these genes at the top of our dataset may merely reflect the sensitive nature of the splicing of genes that are important for neuromuscular biology, and the fact that transcriptome diversity is greatest in neuronal tissues, including the brain [70].

Although we found evidence for both snRNP-dependent and SMN-specific RNA changes in SMA models, it remains to be determined which of these, if any, contributes to disease pathology.

Our analysis of severe and intermediate SMA models suggests that snRNP-independent, *Smn* genespecific RNA changes may play a bigger role in SMA pathology than previously envisioned. The SMN-specific activation of stress signaling pathways is conserved in SMA models [71,72] and correlates well with disease severity in *Smn* hypomorphs (Fig. 8C). This activation of stress signaling appears to be a common signature of neuromuscular disease [73].

Based on our finding of a potential activation in stress signaling in fly models of SMA, an important question moving forward will be to understand how loss of SMN causes this stress response. By text mining with the stress signatures identified in the SMA models described here, we found that mutations in the *Activating transcription factor 3 (Atf3*) cause a similar dysregulation in immune and metabolic homeostasis [74]. Among other things, this signaling pathway appears to regulate the cytosolic formation of SMN-containing U-bodies in human cells in response to bacterial pathogens [75]. If SMN does indeed intersect with a normal host cell response to bacteria, this would provide a parsimonious model for the activation of stress signaling in SMA models, and another avenue to explore potential connections to neuromuscular disease.

In conclusion, a direct comparison of snRNP biogenesis mutants revealed that defects in snRNP supply cause complex changes to the transcriptome. However, an analysis of three different *Smn* hypomorphic lines showed that RNA processing changes are unlikely to be the primary drivers of SMA pathophysiology. Our studies support a model of SMA etiology wherein general disruptions at the mRNA level and specific disruptions at the SMN protein-protein level are responsible for SMA phenotypes. In future experiments, the key will be to uncover the tissue specific molecular activities of SMN that are important for neuromuscular function. Ongoing work with these and other intermediate models of SMA should help clarify the relative contributions of the aforementioned factors to SMA pathology and etiology.

## Materials and methods

### Fly mutants and husbandry

Stocks were cultured on molasses and agar at room temperature (25±1°C) in half-pint bottles. *Oregon-R* was used as the wild-type allele. The *Smn*^*X7*^ microdeletion allele was a gift from S. Artavanis-Tsakonis (Harvard University, Cambridge, USA) [76]. Transposon insertion alleles *Smn*^*D*^ (01109), *Ars2*^*C*^ (*c00735)*, and *Ars2*^*E*^ (*e01128*) were obtained from the Exelixis collection at Harvard Medical School [42,77]. The transposon insertion allele *Phax*^*SH*^ (P{lacW}Phax^SH0641^) was a gift from S. Hou [78].

The wild-type and mutant *Smn* transgenes are expressed using the native *Smn* promoter. Transgenic constructs were integrated into the PhiC31 landing site located at 86F8, which was previously recombined into an *Smn*^*X7*^ background [9,46]. For RNA-seq analysis, *Smn* transgenes in the *Smn*^*X7*^ null background were crossed to *Smn*^*X7*^ animals, and trans-heterozygous animals were put into TRIzol^®^ <12 hours after pupation, as brown prepupa. For qRT-PCR and developmental northerns, *Smn* transgenes in the *Smn*^*X7*^ background were self-crossed, and homozygous animals were put into TRIzol^®^ at 72-77 hours post egg laying or <12 hours after pupation before air bubble migration.

For exogenous expression of a wild-type *Phax* transgene, an intron-less *Phax* cDNA was subcloned into pBID-UASC-GV [79], and subsequently injected and integrated into the 51C1 PhiC31 landing site by BestGene Inc. (Chino Hills, CA). The *UAS:Phax-mVenus* transgene at 51C1 was subsequently recombined into the *Phax*^*SH*^ background. The *armadillo* promoter-GAL4 driver P{GAL4-*SH* arm.S}11 [80] was also recombined into the *Phax*^*SH*^ mutant background.

### RNA-seq & bioinformatics analyses

RNA was isolated from 74 ± 2-hour-old *wild-type* and snRNP biogenesis mutant larvae by homogenization in TRIzol® (Invitrogen^™^), according to the manufacturer’s protocol. Staged animals were isolated from either the wandering third instar stage of development or just before bubble migration in early tan pre-pupae (<12 hours after puparium formation). RNA was DNased with Amplification Grade DNase I (Invitrogen^™^) and TURBO^™^ DNase (Ambion^™^). A TruSeq RNA Sample Preparation Kit v2 (Illumina®) was used for: poly-(A)-enrichment or substituted here with Ribo-Zero^™^ (Epicentre®) rRNA subtraction, barcoding for multiplexing, and cDNA library preparation. Paired end (2x50) sequencing was performed on an Illumina® HiSeq 2000 platform. TopHat and Cufflinks were used for gene expression analyses, according to the bioinformatic pipeline from Trapnell et al. (2012). DaPars was used to identify differences in RNA length and alternative polyadenylation [37,38]. MISO [41] was used to measure changes in annotated alternative-splicing events. For comparison, we used a ‘strong’ Bayes factor cutoff for our poly(A)-selected RNA-seq analysis [81]. We used a ‘positive’ Bayes factor as our cutoff for the total rRNA(-) RNA-seq analysis [82]. For developmental comparison, all publicly available RNA-seq data from L2 to L3 puff stage 7–9 were downloaded from the NCBI Sequence Read Archive (SRA) and converted to fastq format with the SRA toolkit [83] (http://www.modencode.org/). The heatmap was generated with the ‘pheatmap’ package from bioconductor (http://www.bioconductor.org).

### Northern blotting

Northern analyses were performed using standard protocols [84]. Briefly, RNA was isolated as indicated above and approximately 300 ng of total RNA was separated on 10% Novex® TBE Urea gels (Invitrogen^™^), transferred to GeneScreen Plus® charged nylon membranes (PerkinElmer®), dried, and UV cross-linked. Northern blots were probed with 5'-end-labeled oligonucleotides for the respective snRNAs, labeled with T4 polynucleotide kinase (NEB®) and [y-^32^P]ATP (Perkin-Elmer®).

The snRNA-specific oligonucleotide probe sequences are listed in Table S22. Hybridizations were performed at 65°C, and sequential washes in 2x SSC and 0.33x SSC were performed at 60°C. Damp blots were exposed to storage phosphor screens (GE Healthcare) and analyzed with a Typhoon TRIO+ (GE Healthcare). Bands were quantified using ImageQuant TL (GE Healthcare), and *p*-values for three biological replicates were determined with a one-tailed Student’s t-test for unequal variance of two-samples.

RT-PCR analysis Semi-quantitative RT-PCR was performed with intron flanking oligos, using the 2X Apex^™^ Taq Master Mix (Genesee Scientific). Quantitative RT-PCR experiments were carried out as in Praveen et al. (2012). Briefly, random hexamers were used to prime the reverse transcription of isolated RNA according to the manufacturers protocol SupersScript® III (Invitrogen^™^). Real-time PCR reactions of cDNA were carried out on a 7900HT Fast Real-time PCR machine (Applied Biosystems^™^), according to the Maxima® SYBR Green/Rox qPCR Master Mix (2X) (Thermo Scientific^™^) two-step protocol. Oligo pairs amplified cDNAs from intron inclusive versus spliced mature mRNAs. Three biological replicates were tested for each genotype. The ΔΔCt method was used to quantify differences, and *p*-values were determined with a one-tailed Student’s t-test for unequal variance of two-samples. Genespecific primer sequences are listed in Table S22.

### Western blotting

Western analyses were performed using standard protocols and modifications to the methods of Praveen et al. (2012). Briefly, larval protein lysates were prepared by crushing the animals in lysis buffer (50mM Tris-HCl [pH 7.5], 150 mM NaCl, 1mM EDTA, 1% NP-40, 0.1% SDS, 1% Deoxycholate) with 5X protease inhibitor cocktail (Invitrogen^™^) and clearing the lysate by centrifugation. The anti-dUTPase antibody was a generous gift from B.G. Vértessy, and it was used at a dilution of 1 to 50,000 [85]. Affinity purified anti-dSMN was used at a 1 to 2,500 dilution [9]. Antialpha-tubulin antibody (Sigma®) was used at a 1 to 50,000 dilution.

### Cell culture and RNAi

The S2 cell line was obtained from the Drosophila Genome Resource Center (Bloomington, IN) and cultivated according to published protocols [86]. Gene-specific RNAi knockdown was also performed as described in Rogers and Rogers (2008). Briefly, S2 cells were maintained in Sf-900^™^ II SFM media (Gibco^™^) supplemented with 1X Pen Strep (Gibco^™^). For RNAi, one milliliter of (1x10^6^ cells/ml) was aliquoted into each well of a 6-well tissue culture plate. Double-stranded RNA (dsRNA) was *in vitro* transcribed over night (~16 hours) from PCR templates with the T7 MEGAscript kit (Invitrogen^™^), and dsRNA products was subsequently treated with TURBO^™^ DNase (Ambion^™^), according to the respective manufacturers protocols. Oligonucleotide sequences for amplifying the PCR templates are listed in Table S22. PCR templates were amplified from the following plasmids: *Smn*, pAFW-SMN; *Prp6/CG6841*, cDNA clone oFa18909 (GenScript); or *Prp8*, cDNA clone OFa09689 (GenScript). Purified dsRNA was phenol-chloroform extracted, ethanol precipitated, resuspended in RNase-free water (Ambion^™^), and quantified by spectrophotometer. Approximately 15 μg of dsRNA was added to 1×10^6^ cells per each well of a 6-well plate. The dsRNA was added once each day for three successive days. On the fourth day, cells were pelleted and resuspended in TRIzol® (Invitrogen^™^). RNA and proteins were extracted according to the manufacturer’s protocol. Data deposition The data from the *Smn* null animals were deposited in NCBI's Gene Expression Omnibus [87] and are accessible through GEO Series accession number GSE49587 (http://www.ncbi.nlm.nih.gov/geo/query/acc.cgi?acc=GSE49587). The deposition of the additional data discussed here in GEO is pending.

## Acknowledgments

We gratefully acknowledge T.K. Rajendra for his contributions during the early phases of this study. We thank A. Malinová, D. Staněk and S.L. Rogers for their assistance with the S2 cell knockdown experiments, and M.P. Meers for critical reading of the manuscript. We also thank C. Jones and P. Mieczkowski of the UNC High Throughput Sequencing Facility (HTSF). Their helpful discussions on platform choice, sequencing strategy, sample preparation, and sample submission were instrumental to the success of the project. P. Mieczkowski performed the sequencing, and A. Brandt from the HTSF prepared TruSeq cDNA libraries.

## Figure Legends

**Supplemental Figure S1.** Timing of the developmental delays exhibited by the snRNP biogenesis mutants. (A) Pairwise comparisons of Cuffdiff log_10_FPKM levels from age-matched *Oregon-R* and snRNP biogenesis mutants relative to modENCODE developmentally-staged controls for second instar (L2), early 3^rd^ instar (L3 12h), and a later "puff” stage of 3^rd^ instar (L3 P1-2) *wild-type* larvae. Individual points represent individual genes or loci containing multiple genes that were not disambiguated by the Cuffdiff differential expression analysis. (B) Dendrogram comparison of overall gene expression between age-matched *Oregon-R*, snRNP biogenesis mutants, and modENCODE developmentally staged animals. L3 P3-6 and L3 P7-9 are subsequent 3^rd^ instar stages relative to L3 P1-2. The dendrogram is an alternate hierarchical clustering representation of the data presented in Fig. S1A.

**Supplemental Figure S2.** snRNA levels in snRNP biogenesis mutants quantified by developmental northern (A) and ribo-minus total RNA-seq data (B-C). Note that the control flies in panels B and C are age-matched at 72 hrs, and thus RNA-seq reads are artificially high in the controls. (B) snRNA levels from aggregated mapped read counts, normalized to U6 snRNA mapped read counts. (C) snRNA levels from aggregated mapped read counts that were normalized to the median of the middle two quartiles of all mapped *WT* sequence read counts. Normalized age-matched *Oregon-R* (*WT*) snRNA levels were set at 100.

**Supplemental Figure S3.** Relative abundance of snRNA isoforms in snRNP biogenesis mutants: (A) U1 snRNA paralogs; (B) U2 snRNA paralogs; (C) U4 snRNA paralogs; and (D) U5 snRNA paralogs. Identical isoforms are listed together.

**Supplemental Figure S4.** Normal developmental changes in mRNA length. Pairwise comparison of significant differences in mRNA length between modENCODE developmentally staged animals: (A) L2 and L3 12h; (B) L3 12h and L3 P1-2; and (C) L2 and L3 P1-2.

**Supplemental Figure S5.** Semi-quantitative RT-PCR of alternative-splicing changes identified by RNA-seq. MISO alternative-splicing event annotations are listed.

**Supplemental Figure S6.** Analysis of snRNA levels in transgenic rescue lines. (A) Northern blot of snRNA in the *Phax* mutant and *Phax* rescue line (*Arm>Phax*) relative to *WT*60h larvae; quantified in panel (B). (C) Northern blot of snRNA levels in the *Smn* mutant and *Smn* rescue line (Tg:*Smn*^*WT*^) relative to *WT* 60h; quantified in panel (D). U6-normalized snRNA levels of *WT* were set at 100.

**Supplemental Figure S7.** snRNP-specific protein knockdown in S2 cells. (A) qRT-PCR of untreated cells versus those treated with dsRNA for *Smn, Prp6* or *Prp8* mRNAs. (B) Western blot of dUTPase levels in RNAi-treated cells. Anti-SMN and anti-alpha-tubulin verify SMN knockdown and load, respectively. (C) qRT-PCR verification of *Prp6* and *Prp8* mRNA knockdowns. Levels in untreated cells were set at 100. (D) Northern blot of snRNA levels from S2 cell knockdowns, quantified in (E). U6-normalized snRNA levels of *WT* were set at 100.

**Supplemental Figure S8.** Lack of snRNA level decreases in *Smn* missense mutants over development. (A&C) Northern blots of snRNA levels in the *SMN* missense mutants over development, quantified in (B&D). snRNA levels from the *Smn*^*WT*^ (WT) transgenic lines were set at 100. Pupal animals were staged by color and internal air bubble, and larval animals were from a timed collection (72-77 hours post egg laying). The small differences in the early larval stage likely reflect differences in developmental timing, as the *Smn* missense mutants exhibit slower developmental progression.

**Supplemental Figure S9.** Intron retention in *scarface* transcript. (A) qRT-PCR analysis of intron retention of flies expressing SMA patient-derived missense mutations. Transgenic animals of the following generalized genotype were used: *Smn*^*X7/X7*^*,Flag-Smn*^*Tg/–*^, where Tg represents a WT, V72G, Y107C, or T205I transgene. Intron retention in the *Smn*^*WT*^ transgenic rescue line was set at 1. (B) qRT-PCR of untreated cells versus cells treated with dsRNA for *Smn* or *Prp8* mRNAs.

